# Experience- and sex-dependent intrinsic plasticity in the zebra finch auditory cortex during song memorization

**DOI:** 10.1101/658104

**Authors:** Andrew N Chen, C Daniel Meliza

## Abstract

Early auditory experience is critical to the development of vocal communication. Zebra finches and other songbirds have a sensitive period when young birds memorize a song to use as a model for vocal production. We found that intrinsic spiking dynamics change dramatically during this period in the caudal mesopallium, a cortical-level auditory area. Specifically, the proportion of neurons that only fire transiently at the onset of intracellular current injections increases, along with Kv1.1, a channel that facilitates transient spiking. Plasticity is greater in males and requires exposure to a complex, noisy environment. These observations indicate that intrinsic dynamics are modulated in response to the acoustic environment to support robust auditory processing during a critical phase of vocal learning.

## Main Text

Vocal communicators require auditory systems that can reliably encode information over the broad range of spectral and temporal scales found in vocalizations (Singh and Theunissen, 2003; Wang et al., 2008). In humans, songbirds, and other species that learn how to vocalize through sensorimotor learning, the auditory system must also provide reliable feedback so that motor production can be adjusted to match desired targets. Early experience hearing conspecific vocalizations not only determines the targets to be imitated, but also influences auditory perception (Doupe and Kuhl, 1999). A classic example is the perceptual narrowing that occurs in humans during the first year of life, in which exposure to speech establishes the phonetic distinctions an infant is able to perceive (Werker and Lalonde, 1988; Kuhl et al., 1992; Kuhl, 2004). The neural plasticity underlying this critical stage in development is poorly understood.

As in humans, early auditory experience is critical to vocal communication in zebra finches (*Taeniopygia guttata*). Around 25 days of age, juvenile males begin to memorize the song of an adult male tutor whom they will learn to imitate (Gobes et al., 2017). Birds who do not hear song during this sensory acquisition period are unable to produce species-typical songs as adults, even if they are exposed to a tutor later in life (Marler and Tamura, 1964; Eales, 1985). To understand this process and relate it to human speech development, it is important to consider not only how specific song memories guide sensorimotor learning, but also how the early acoustic environment (including but not limited to the song of a primary tutor) might shape auditory processing more generally (Woolley, 2012). In support of this idea, raising birds in acoustically impoverished environments produces a range of behavioral and physiological deficits consistent with abnormal auditory processing (Sturdy et al., 2001; Chen et al., 2017; Amin et al., 2013). Conversely, birds raised without exposure to a tutor but in an acoustically rich environment retain some ability to learn from a tutor later in life (Morrison and Nottebohm, 1993), suggesting that auditory processing is spared enough to allow learning to occur. Thus, auditory development supports but is dissociable from song memorization.

We set out to determine the neural basis underlying the effects of early experience on auditory processing. This study focuses on the caudal mesopallium (CM), a cortical-level auditory area (Wang et al., 2010) that responds selectively to familiar conspecific songs (Gentner and Margoliash, 2003; Jeanne et al., 2011; Meliza and Margoliash, 2012) and could therefore be a site of plasticity during development. In juveniles, the putatively excitatory neurons in CM have diverse intrinsic spiking dynamics. Some neurons produce sustained responses to depolarizing step currents, while others respond only transiently at the beginning of the stimulus (i.e., phasic excitability). We previously showed that phasic excitability is associated with a low-threshold potassium current, that it improves encoding of rapid modulations at the frequencies found in zebra finch song, and that it can make neural responses to song less sensitive to noise (Chen and Meliza, 2018; Bjoring and Meliza, 2019). These findings suggest that phasic excitability in CM supports robust auditory processing. Thus, we asked here whether intrinsic excitability among CM neurons changes during development in response to early experience.

We measured intrinsic excitability in zebra finches starting at age P18, when chicks fledge from the nest, continuing through the sensory acquisition and sensorimotor learning periods, and into adulthood (Fig. 1A). We made whole-cell recordings from slices of CM (Fig. 1B) and characterized the spiking patterns produced in response to step currents. As in our previous study, the putatively excitatory (broad-spiking) neurons exhibited diverse patterns of spiking responses, ranging from neurons that produced sustained responses throughout the stimulus (i.e., tonic excitability; Fig. 1C) to neurons that only fired a single spike at the onset (i.e, phasic excitability; Fig. 1D). We quantified phasic excitability using a continuous metric that was simply the log-scaled response duration averaged across all current steps (∆*t*); decreases in response duration indicate a greater tendency for phasic firing. We also classified each neuron as tonic or phasic, depending on whether it responded in the last 500 ms of the stimulus for any current stimulus.

**Fig. 1.**
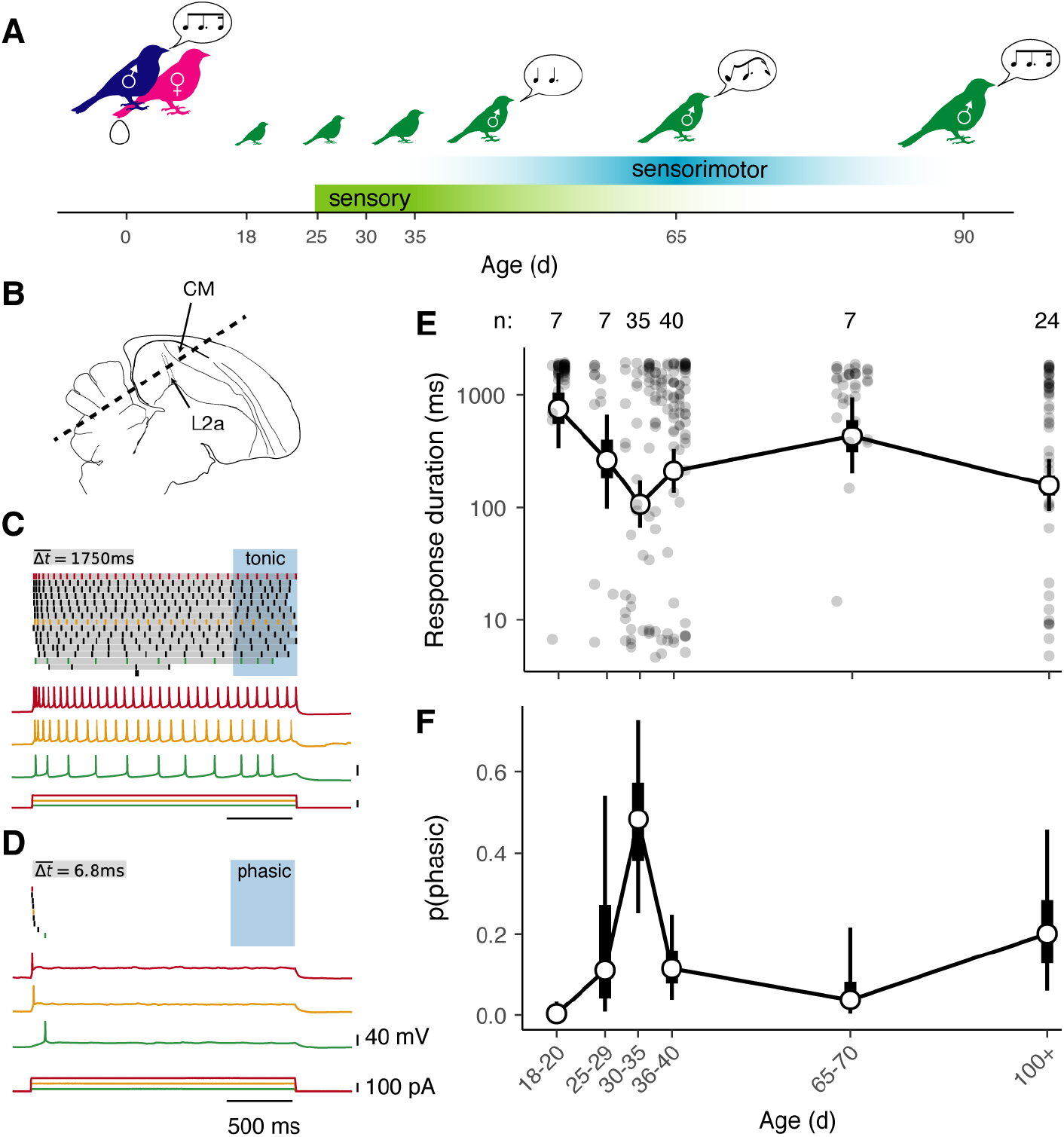
Intrinsic dynamics of CM neurons change over development. (**A**) Diagram of zebra finch auditory and vocal learning. The sensory learning phase, when birds memorize a song to copy, overlaps with a later period of sensorimotor learning. (**B**) Parasaggital diagram of the zebra finch brain with the caudal mesopallium (CM) indicated in relation to the primary auditory thalamorecpient area L2a. Dashed line indicates the orientation of the brain slices cut for whole-cell recordings. (**C**) Example of tonic spiking from a P20 male. Top is a spike raster plot showing responses to 2 s current steps of increasing amplitude. The grey bars illustrate how average response duration (∆*t*) is calculated, and the blue-shaded region shows the last 500 ms of the response. Because the neuron spiked at least once during this window, it was classified as tonic (see materials and methods). Below are voltage recordings and current steps from three selected trials, with colors corresponding to trials in the raster plot. (**D**) Example of phasic spiking from a P37 bird of unknown sex. The neuron fired a single spike to all current steps. (**E**) Average response duration as a function of age. Solid points correspond to individual neurons. Hollow circles show means for age ranges indicated by labels on horizontal axis. Thick and thin whiskers show 50% and 90% credible intervals for the mean (GLMM; see materials and methods). Numerals above the plot indicate the number of birds in each group. Compared to P18–P20, response duration is significantly lower (more phasic) at P30–P35 (p = 0.002), P36–P40 (p = 0.014), and in adults (p = 0.010). Additionally, compared to P30–P35, average response duration is significantly greater (less phasic) at P36–P40 (p = 0.038) and P65–P70 d (p = 0.002). (**F**) Estimated proportion of neurons that are phasic as a function of age (GLMM). Compared to P18–P20, phasic neurons are more prevalent at P30–P35 (p < 0.001), P36–40 (p = 0.018), and in adults (p = 0.006). Compared to P30–P35, phasic neurons are less prevalent at P36–P40 (p = 0.012) and P65–P70 (p = 0.020). Group sizes are as indicated in **E**.

Although a wide variety of excitability patterns were present among CM neurons from birds across the entire age range studied, the distribution of intrinsic dynamics differed dramatically across development. In newly fledged birds (P18–P20), almost every neuron was tonic. Phasic excitability increased with age, reaching a peak between P30–P35 (Fig. 1, E and F). After this peak, phasic firing decreased as the sensory acquisition phase closed (P65–P70) and then increased again in adulthood (P100+).

Because the peak in phasic excitability occurred at the height of sensory acquisition for song, we hypothesized that this intrinsic plasticity was involved in song memorization, either directly by supporting the encoding of a specific memory, or perhaps more indirectly through a permissive role that facilitates auditory discrimination. To test this idea, we tracked changes in intrinsic firing patterns among CM neurons over time in birds that were not exposed to song (i.e., chicks housed in acoustic isolation boxes with only their mothers; Fig. 2A). In contrast to colony-reared (CR) controls, phasic firing was essentially absent in these female-reared (FR) animals during sensory acquisition (Fig. 2, B to D), though it appeared in older animals (P65–70 and adults). Thus, intrinsic dynamics are affected by experience, but primarily during song memorization.

**Fig. 2.**
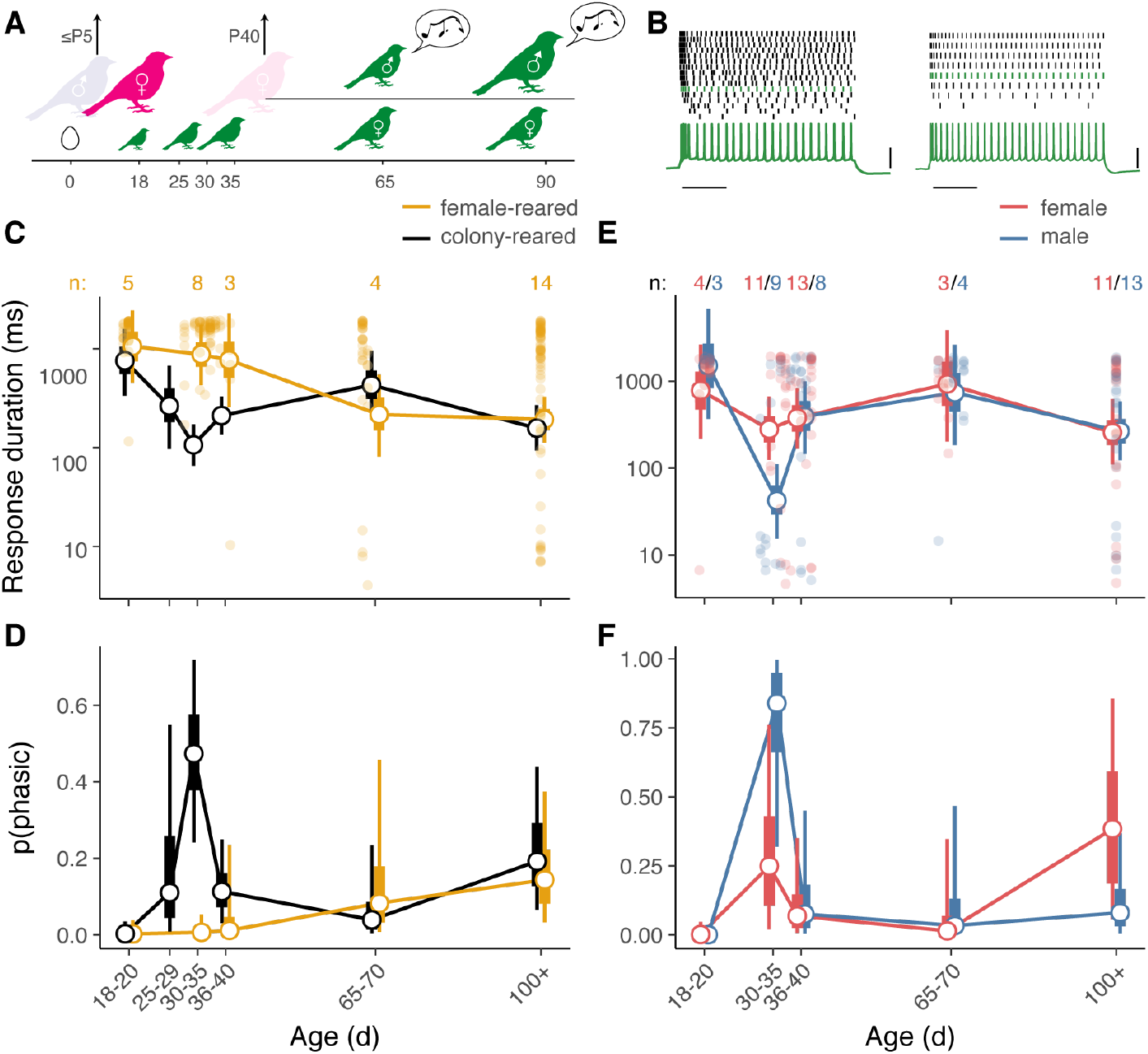
Phasic firing during the sensory learning period depends on sex and experience. (**A**) Diagram of female-rearing paradigm. The family is placed in acoustic isolation and the father removed 1–5 d after the first egg hatches. The mother is removed around P40 and the male and female chicks are separated. Male chicks raised in this condition produce abnormal (isolate) song. (**B**) Examples of responses to step currents obtained from female-reared chicks (P20 and P32), as shown in Fig. 1C,D. Scale bars, 500 ms and 40 mV.(**C**) Average response duration as a function of age in female-reared (FR; yellow) and colony-reared (CR; black) animals. The format of the plot is as in Fig. 1E. For visual clarity, data points for individual neurons are only shown for female-reared birds. Numerals indicate the number of female-reared birds. Between P30–P35, response duration is lower (more phasic) in CR vs FR birds (p < 0.001). Among FR birds, average response duration at P18–P20 is significantly greater (less phasic) than at P65–P70 (p = 0.04) and in adults (p = 0.002). (**D**) Estimated proportion of phasic neurons. Phasic neurons are significantly more prevalent in CR vs FR animals at P30–P35 (p < 0.001), and among FR animals, more prevalent in adults than at P18–P20 (p = 0.026). (**E**) Average response duration as a function of age and sex in CR animals. Responses are on average shorter in males compared to females between P30–P35 (p = 0.004). (**F**) Proportion of phasic neurons as a function of age and sex. The apparent difference between males and females at P30–P35 is not statistically significant (p = 0.16).

The presence of sex-related differences in intrinsic excitability would also point to a relationship between intrinsic plasticity and song memorization, as only male zebra finches sing. To test this idea, we compared male and female CR birds. Plumage becomes sexually dimorphic in zebra finches around P40; in younger birds, we were able to determine the sex of a subset of animals (n = 48/89) by genotyping DNA from blood or feathers. As predicted, intrinsic dynamics were more phasic in males compared to females aged P30–P35, i.e., when plasticity is experience-dependent and phasic firing is most prevalent (Fig. 2, E and F). The difference between males and females was statistically significant for response duration but not for the proportion of phasic neurons (Fig. 2E), indicating that the effect of sex is weaker than that of experience during P30–P35. In adults, there was no difference in phasic excitability between males and females.

We also compared older male and female FR birds to determine if the later phase of intrinsic plasticity depended on experience. Male FR birds go through sensorimotor vocal development despite the absence of a model to copy (Konishi, 1965; Volman and Khanna, 1995). Although the learning process converges on aberrant “isolate song” in these birds, they do still vocalize, and thus could drive experience-dependent plasticity in themselves and among nestmates. Therefore, we separated male and female FR chicks around P40 (the age at which sex is apparent based on plumage differences), thereby minimizing the exposure of young females to the attempts of their male siblings to sing. There was a trend towards more phasic excitability among CM neurons in adult FR males compared to adult FR females (Fig. S1), but it was not statistically significant (GLMM: p = 0.096; n = 12 birds). These data suggest that the later increase in phasic excitability is largely independent of experience.

We next sought to identify the specific acoustic factors required for intrinsic plasticity to occur during the early, experience-dependent period. Although FR birds heard the calls of their mother and siblings, they were not exposed to normal adult song, which is exclusively produced by males in this species. The isolation boxes also create a much quieter, less complex acoustic environment for the FR birds. In the CR condition, chicks lived in individual cages with their father, mother, and siblings, and the cages were situated in a room containing 5–10 other pairs and families. These conditions mimic the rich acoustic environment of colony noise–songs and calls from many different individuals–that these highly social birds would experience in the wild. Colony noise is characterized by large amplitude modulations and a complex spectrotemporal modulation spectrum (Fig. S2).

To dissociate the effects of specific types of early auditory exposure, i.e., tutor song versus colony noise, we examined two additional rearing conditions (Fig. 3A). Like FR birds, pair-reared (PR) chicks were housed in acoustic isolation boxes. However, under the PR condition the father was not removed. Thus, PR chicks heard song in a quiet environment and receive the same level of parental care and social interaction as CR chicks. Visual isolates (VI) were housed in the colony room, but in the absence of the father and with the addition of opaque barriers between cages. Thus, VI chicks heard colony noise, but could not socially interact with any adult males. This lack of social interaction appears to be critical for song acquisition, as despite hearing song, VI chicks produce abnormal adult songs that resemble those of FR birds (Morrison and Nottebohm, 1993). After VI and PR chicks reached 30–40 days of age, we recorded responses to step currents in CM slices. (This experiment was carried out before we had fully identified the narrow period between 30–35 d when phasic excitability is at its peak, so a broader range of ages was used). Surprisingly, the effect of exposing birds to a tutor (the PR condition) was small and not statistically significant, whereas exposing birds to colony noise in the absence of a tutor (the VI condition) partially rescued phasic excitability. The effect of VI rearing nearly reached statistical significance (Fig. 3, B and C), despite the small sample size (n = 4 birds). This result suggests that a complex acoustic environment, rather than song exposure, is necessary for phasic firing to emerge.

**Fig. 3.**
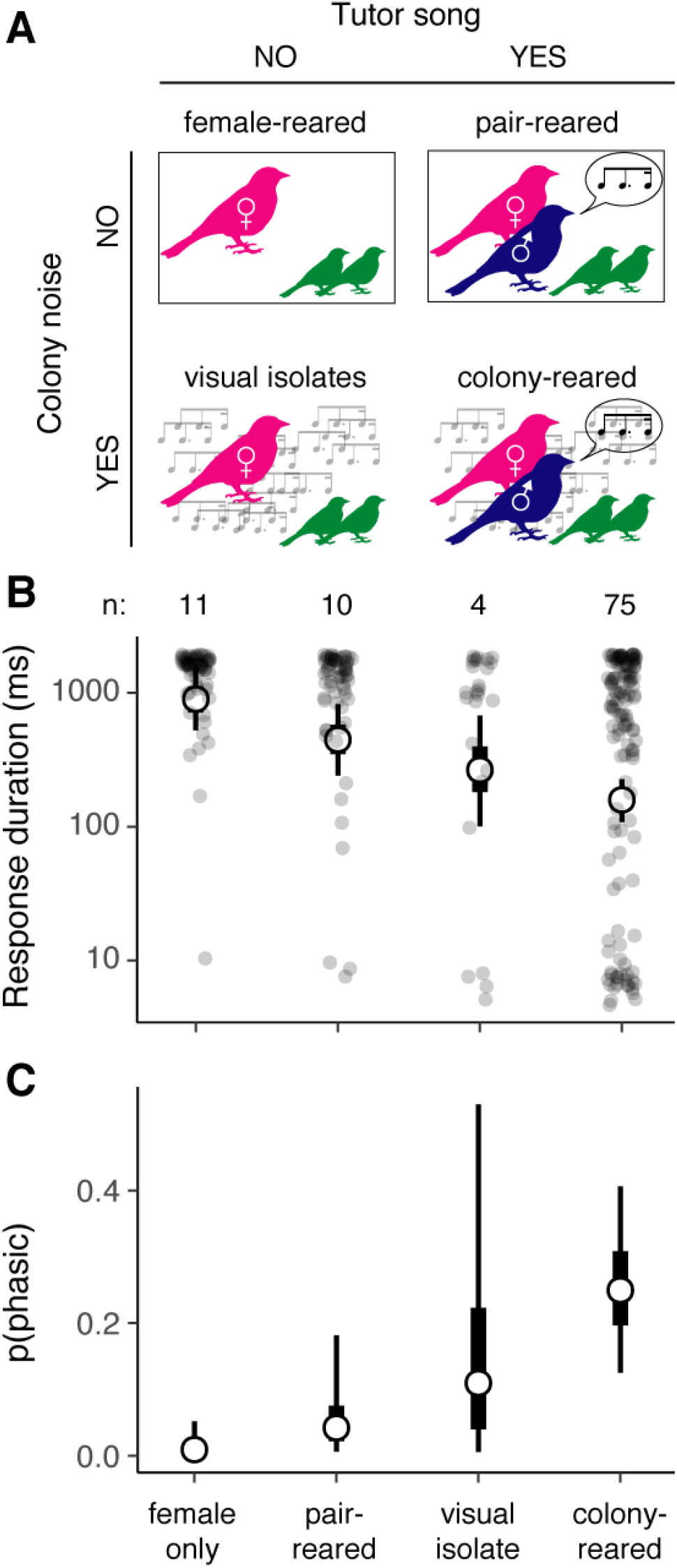
Phasic dynamics are partially rescued by exposure to a complex acoustic environment. (**A**) Schematic of rearing conditions. Only pair-reared and colony-reared birds have direct social interaction with a male tutor, whereas only visual isolates and colony-reared birds are exposed to noise comprising songs, calls, and other sounds from the colony (Fig. S2). (**B**) Average response duration in P30–P40 birds for each condition. Format is as in Fig. 1E. Compared to female-reared birds, average response duration is significantly lower (more phasic) in colony-reared birds (GLMM: p < 0.001) and nearly significantly lower in visual isolates (p = 0.052). (**C**) Proportion of phasic neurons in each condition. Compared to female-reared birds, there are more phasic neurons in colony-reared birds (p = 0.006).

Rather than add more animals in pursuit of statistical significance, we confirmed the result by directly testing the hypothesized mediating mechanism, i.e., changes in Kv1.1 expression. Kv1.1 is a voltage-gated potassium channel that activates at low voltages (Bal and Oertel, 2001), and which is responsible for phasic firing in several other auditory areas (Rothman and Manis, 2003; Sivaramakrishnan and Oliver, 2001). The gene for Kv1.1, KCNA1, is transcribed in zebra finch CM (ZEBrA Database, 2019), but it is not known whether protein levels change over the course of development and/or with experience. Using a monoclonal antibody against the C-terminus of rat Kv1.1, we found that Kv1.1 was highly expressed in CM neurons during P30–P35 (Fig. 4A). The proportion of Kv1.1-expressing neurons was higher in P30–P35 CR birds compared to age-matched FR animals and younger CR birds (Fig. 4B). In VI birds, there were also more Kv1.1-expressing neurons than in age-matched FR animals, though somewhat less than in CR birds, consistent with our electrophysiological results (Fig. 3, B and C). In all rearing conditions, staining appeared punctate and was primarily located in the cell body outside the nucleus, or occasionally distributed along what looked like processes (Fig. 4C). The proportion of Kv1.1 neurons was consistently higher in more medial sections (odds ratio = 1.19; p = 0.016), but there was no difference between sexes (p = 0.70) or between hemispheres (p = 0.88). There is no well-validated membrane marker in zebra finches, so we were unable to determine what proportion of the signal was colocalized with the plasma membrane.

**Fig. 4.**
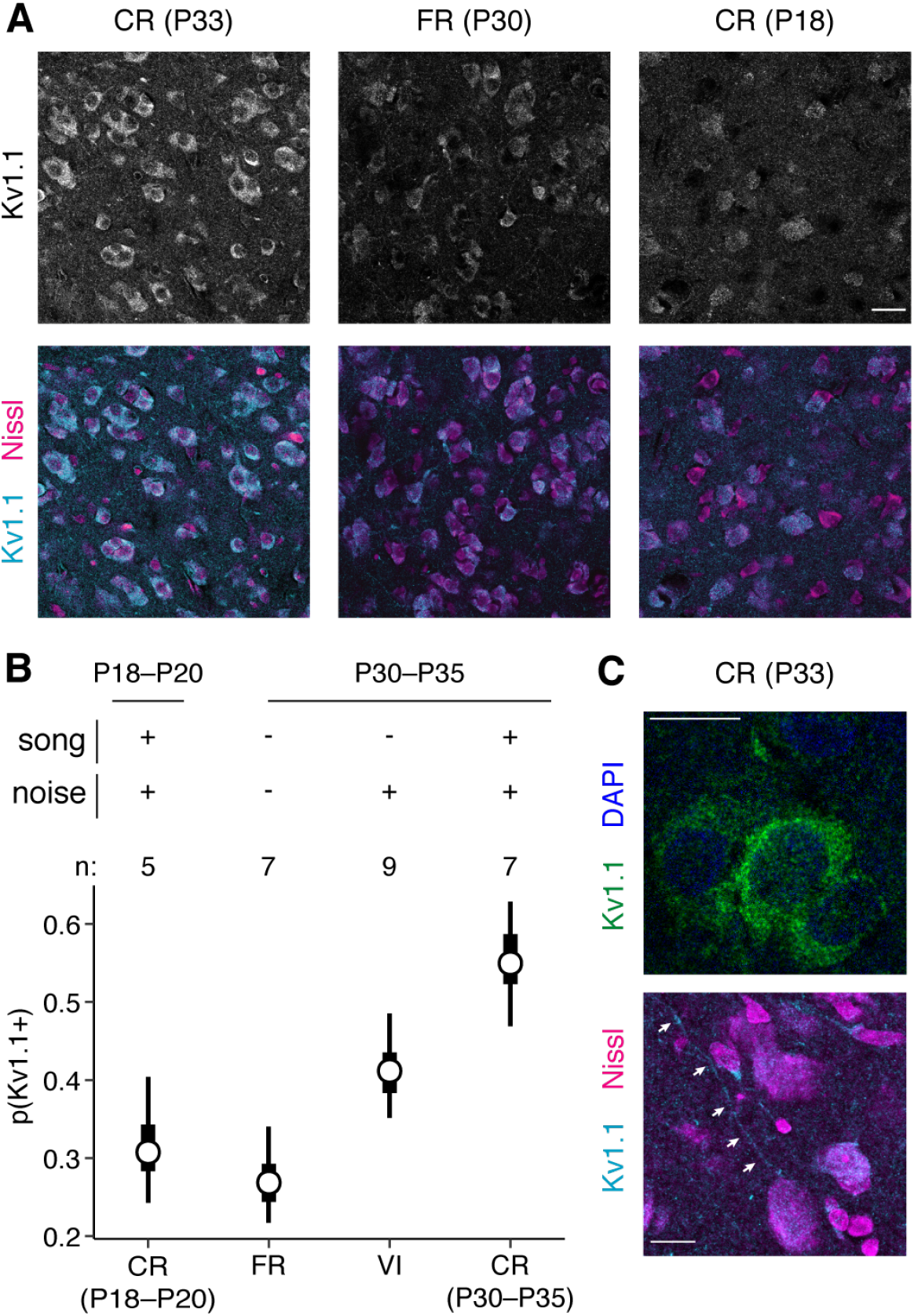
Kv1.1 expression depends on age and experience. (A) Examples of *α*-Kv1.1 staining in different rearing conditions. Images are single confocal slices (0.44 µm) with identical laser, gain, and contrast settings for Kv1.1. Columns correspond to samples from different ages and rearing conditions. Top row shows Kv1.1 signal in grayscale; bottom row is a composite of Kv1.1 (cyan) with a fluorescent Nissl stain (magenta). Scale bar, 20 µm. (B) Estimated proportion of Kv1.1-positive neurons at each age/condition tested. Compared to FR birds at P30–P35, the proportion of Kv1.1-positive neurons is higher in age-matched CR animals (GLMM: p < 0.001) and VI animals (p = 0.01). Numerals indicate the number of animals in each group. (C) Higher-power images (63X) of Kv1.1 staining in cell bodies (top) and processes (bottom). Top image shows composite of Kv1.1 (green) with DAPI (blue); bottom shows composite of Kv1.1 (cyan) with fluorescent Nissl (magenta). Scale bars, 10 µm.

Our results demonstrate that intrinsic neuronal dynamics in CM are highly plastic during development. Between fledging and adulthood, there are three shifts in the distribution of intrinsic dynamics, and the timing of these changes coincides with key milestones in the development of vocal communication in this species. In the youngest animals we examined (P18–P20), the overwhelming majority of broad-spiking neurons exhibited sustained, regular spiking patterns typical of excitatory cortical neurons. At the age when young males begin to memorize a song to copy (P30–P35), approximately half of the cells had shifted to only producing transient responses. Within several days (P36–P40), intrinsic dynamics then shifted back to tonic firing. By the time males had started intensively practicing song production (P65–P70), only about 10% of the CM neurons we recorded from were were phasic. Phasic excitability again increased in adults, i.e., at ages when males have finalized the song they will produce for the rest of their lives, and when both sexes are employing song in social interactions (including, but not limited to mate selection). Each of these phases of plasticity depend on experience and sex to differing degrees.

The earliest phase of intrinsic plasticity, which occurs at the beginning of the song memorization period for young male zebra finches, is dependent on experience and sex. We find that phasic excitability fails to emerge during this period in FR birds, and is stronger in males than in females. However, we also find that hearing song from a tutor is not sufficient to drive intrinsic plasticity; instead, phasic excitability requires exposure to a noisy acoustic environment. This suggests that the intrinsic plasticity occurring during this earliest phase is not directly involved in storing memories of a song to be copied, but instead is an adaptation to the acoustic environment in which memorization is occurring. We discuss this idea in more detail below, after considering the mechanism and the other phases of plasticity.

The dramatic change in physiological properties that occurs during song memorization is mirrored by increased expression of Kv1.1, a low-threshold potassium channel that facilitates phasic firing. Manipulating the acoustic environment produces similar effects on both electrophysiological properties and Kv1.1 expression, i.e., both shift towards more phasic-like conditions. Among CR birds aged P30–P35, there is close quantitative agreement between the proportion of Kv1.1-expressing neurons and the proportion of phasic neurons identified through electrophysiology, indicating that our whole-cell recordings represent a relatively unbiased sample of the population. However, the correlation is not as good in birds raised under other conditions: Kv1.1 is expressed in close to 30% of cells in FR and younger CR animals, but there are essentially no electrophysiologically phasic neurons in either of these groups. This discrepancy likely stems from two different sources. First, the effect of low-threshold potassium currents on firing patterns is highly nonlinear, and a minimum level is required to effectively suppress repetitive firing (Rothman and Manis, 2003; Chen and Meliza, 2018). Second, much of the Kv1.1 staining we observed was punctate and within the cell body (Fig. 4C), suggesting that a large fraction of the channels are within the cell, not at the cellular membrane where they need to be to influence intrinsic dynamics. Indeed, Kv1.1 has a strong ER retention signal (Manganas et al., 2001; Vacher et al., 2007), so expression is likely necessary, but not sufficient for phasic excitability. This result is consistent with activity- and experience-dependent regulation of Kv1.1 expression and localization reported in a number of other systems (Lu et al., 2004; Dehorter et al., 2015; Akter et al., 2018).

Experience-dependent plasticity in sensory systems is common during development (Sanes and Woolley, 2011; Levelt and Hübener, 2012). In rodent auditory cortex, manipulating experience during an early critical period has long-lasting effects on functional response properties, (Zhang et al., 2001; de Villers-Sidani et al., 2007; Zhou and Merzenich, 2008), perception (Han et al., 2007; Kover et al., 2013), synaptic connections (Dorrn et al., 2010; Sun et al., 2010) and intrinsic properties (Kotak et al., 2005). In zebra finches, experience hearing tutor song affects activity-dependent gene expression (Gobes et al., 2010), perception (Sturdy et al., 2001; Chen et al., 2017), functional response properties (Adret et al., 2012; Yanagihara and Yazaki-Sugiyama, 2016), and intrinsic excitability (Ross et al., 2019). The early, experience-dependent plasticity we observed is consistent with these findings and with the general consensus that sensory circuits adapt during development to the structure of sensory input. Indeed, whereas many previous studies required deafening or thousands of repetitions of artificial stimuli to induce plasticity, we found that natural noise, only 15 dB louder than the FR condition, was sufficient to cause large changes in channel expression and physiology. Our results therefore provide a new level of support for the biological significance of developmental plasticity to auditory perception (Bao et al., 2013).

In the second phase of intrinsic plasticity, phasic excitability decreases almost to the P18–P20 baseline. Although we sampled at only a few older ages, the timing of the decrease is suggestive, as P65 is around the time when the sensory acquisition phase ends, and birds are no longer able to learn new song elements (Gobes et al., 2017). If, as we posit, phasic excitability facilitates both auditory processing development and song learning, then the shift back to more tonic dynamics at this age could contribute more directly to the process of song acquisition; specifically, a diminished ability to learn new material during sensorimotor learning may be regulated by the processes that cause the song memorization period to close (London, 2017; Kelly et al., 2018). This hypothesis could be tested by determining if phasic excitability remains high in VI animals, who retain the ability to learn new song material into adulthood (Morrison and Nottebohm, 1993).

The third phase of plasticity occurs as birds become adults. Phasic excitability again increases, but in contrast to the early phase, this change does not appear to depend on experience or sex, as there were no statistically significant differences in adults between FR and CR animals, or between males and females. This result suggests that phasic dynamics have a general function in auditory perception that extends beyond song memorization. As adults, both male and female zebra finches use song to communicate, either as sender and receiver (males) or only as a receiver (females). Thus, both sexes share similar requirements for robust and sensitive auditory perception, which could explain why the late phase of plasticity does not depend on sex.

What is the function of phasic excitability in auditory perception? Although we did not directly address this question in the present study, our observation that intrinsic plasticity is driven by exposure to noise suggests that it is an adaptation to noise. As vocal learners raised in social environments, zebra finch chicks are faced with a challenging version of the cocktail-party problem (Cherry, 1953): not only do they have to separate the song of a single tutor from a background with nearly identical spectrotemporal statistics, they presumably are still learning the structure and nuances of their species’ song. The phasic dynamics generated by Kv1.1 may be critical to solving this problem, especially in younger animals in whom inhibitory circuitry may still be developing (Takesian and Hensch, 2013; Vallentin et al., 2016). In subcortical auditory areas, low-threshold potassium currents facilitate detecting brief periods of coherent excitation within a noisy signal (Golding et al., 1995; Svirskis et al., 2002; Meng et al., 2012), an effect that we also observed in a recent computational modelling study of CM (Bjoring and Meliza, 2019). Although birds raised in quiet environments show no overt deficits in learning from a tutor under similar conditions (Tchernichovski and Nottebohm, 1998), these results suggest that they may be less able to learn if tutoring takes place in a noisier, more natural environment. Similarly, we expect that blocking phasic excitability in adults would negatively impact the ability of both sexes to discriminate songs in noisy conditions (Narayan et al., 2007; Schneider and Woolley, 2013).

Humans are also vocal learners. And like zebra finches, human infants also have an early critical period during which they have to learn the structure of conspecific vocalizations, specifically the acoustic-phonetic mappings of their native language (Werker and Lalonde, 1988; Maye et al., 2002; Kuhl, 2004). Deficits in this learning process may contribute to a number of prevalent speech and language disorders, including dyslexia, that have been linked to poor phonological processing (Mody et al., 1997; Tsao et al., 2004; Ziegler et al., 2005; Pennington and Bishop, 2009; Goswami et al., 2011). Genes implicated as risk factors for these disorders, including CNTNAP2 and KIAA0319, are involved in Kv1.1 trafficking (Strauss et al., 2006) and in regulation of intrinsic excitability (Centanni et al., 2014). Although phasic spiking appears to be rare in mammalian auditory cortex (Metherate and Aramakis, 1999), Kv1.1 is widely expressed in excitatory and inhibitory cortical neurons (Higgs and Spain, 2011; Dehorter et al., 2015), so experience-dependent intrinsic plasticity may also have broad implications for verbal communication in humans. Thus, we recognize that the mechanism implicated in our data may be relevant to future studies of the pathogenesis and therapeutic interventions related to these conditions.

## Materials and Methods

### Experimental Design

The objective of the study was to determine how the intrinsic spiking dynamics of neurons in the caudal mesopallium depend on age, sex, and experience. Ages were sampled between P18 (i.e., 18 d post hatch) and P344, then binned into intervals corresponding to key developmental stages. Experience was manipulated along two dimensions: presence of the father and exposure to the acoustic environment of the colony. The main contrast in the study was between colony-reared (CR) birds, who were housed with both parents in a cage placed in our breeding colony room, and female-reared (FR) birds, who were housed without their father in an acoustic isolation box. Pair-reared (PR) birds were housed in acoustic isolation with both parents. Visually isolated (VI) birds were housed in the colony room but were prevented from visually interacting with their father or any other male. The effects of age, sex, and acoustic experience were measured using whole-cell electrophysiology and immunohistochemistry.

### Animals

All procedures were performed according to NIH guidelines and protocols approved by the University of Virginia IACUC. Zebra finches (*Taeniopygia guttata*) were bred from our local colony. All birds received finch seed (Abba Products, Hillside NJ) and water ad libitum and were kept on a 16:8 h light:dark schedule in temperature- and humidity-controlled rooms 22–24 °C.

Sex was determined from plumage coloration in older animals or by PCR amplification of the CHD-1 gene (Jensen et al., 2003), which has different lengths on the Z and W sex chromosomes. Genomic DNA was isolated from feathers (Qiagen DNAeasy Blood and Tissue Kit) or from blood stored in a 5% Chelex-100 solution. The forward primer sequence was YTKCCAAGRATGAGAAACTG, and the reverse primer sequence was TCTGCATCACTAAAKCCTTT. Because females are heterogametic, these primers yield two products approximately 350 and 400 bp in length in females, whereas male DNA only yields the shorter product. No-template and positive male and female controls were included in each PCR batch. Note that any contamination would result in males being misidentified as female, which would reduce the size of the sex difference observed here rather than producing a false positive.

### Experimental Rearing Conditions

All of the animals in the study were bred in individual cages that were initially placed in a room housing dozens of male and female finches of varying ages. CR chicks remained in the colony room until they were used in an experiment. Juveniles were separated from their parents after about 35 d and housed with siblings or in large aviaries. In the FR condition, the cage was moved into an acoustic isolation box (Eckel Industries, Cambridge, MA) after the first egg hatched. Within 5 d, the father was removed from the cage and housed elsewhere. In some cases, an unrelated female was added to assist with provisioning. Male and female FR chicks were separated after they began to show plumage differences (around 40 d) so that females would not be exposed to FR males’ attempts to sing. PR families were also moved to acoustic isolation boxes after the first chick hatched, but the father was not removed. VI chicks remained in the colony room, but in cages that were separated from each other by opaque barriers. As with the FR condition, the father was removed within 5 d of the first egg hatching. Visual isolates could therefore hear the songs of adult males and call to them but could not socially interact. Some subjects may have been able to see males in a large flight aviary over 1 m away.

### Brain Slice Preparation and Electrophysiology

Acute brain slices were prepared using methods from Chen and Meliza (2018). Zebra finches were administered a lethal intramuscular injection of Euthasol (pentobarbitol sodium and phenytoin sodium; 200 mg/kg; Hospira) and perfused transcardially with ice-cold cutting buffer (in mM: 92 NaCl, 2.5 KCl, 1.2 NaH2PO4, 30 NaHCO3, 20 HEPES, 25 glucose, 5 sodium ascorbate, 2 thiourea, 3 sodium pyruvate, 10 MgSO4⋅7H2O, 0.5 CaCl2⋅2H2O; pH 7.3–7.4, 300–310 mmol/kg). In older animals, the cutting buffer contained 93 mM N-methyl-D-glucamine (NMDG) instead of NaCl (Zhao et al., 2011). The brain was blocked using a custom 3D printed brain holder (http://3dprint.nih.gov/discover/3dpx-003953). 300 µm sections were cut in room-temperature cutting buffer on a VF-200 Compresstome (Precisionary Instruments), and then transferred to 32 °C cutting buffer for 10–15 min for initial recovery. Sections were then transferred to room-temperature holding buffer (in mM: 92 NaCl, 2.5 KCl, 1.2 NaH2PO4, 30 NaHCO3, 20 HEPES, 25 glucose, 5 sodium ascorbate, 2 thiourea, 3 sodium pyruvate, 2 MgSO4⋅7H2O, 2 CaCl2⋅2H2O; pH 7.3-7.4, 300–310 mmol/kg) to recover for at least 1 h before use in recordings. All solutions were bubbled continuously with 95% O2 ⋅ 5% CO2 mixture starting at least 10 minutes prior to use.

Slice recordings were conducted in a RC-26G recording chamber (Warner Instruments) perfused with standard recording ACSF (in mM: 124 NaCl, 2.5 KCl, 1.2 NaH2PO4, 24 NaHCO3, 5 HEPES, 12.5 glucose, 2 MgSO4⋅7H2O, 2 CaCl2⋅2H2O; pH 7.3–7.4, 300–310 mmol/kg) at a rate of 1–2 mL/min at 32 °C, which was the highest and closest to physiological temperatures at which slices remained healthy and recordings stable. Whole-cell patch-clamp recordings were obtained under 60X infrared (900 nm) DIC optics. CM was located relative to LMV and the internal occipital capsule (CIO), which both comprise dense myelinated fibers visible as dark bands under brightfield or IR illumination. Most neurons were recorded from the lateral subdivision of CM (CLM), which is superior to CIO in this sectioning plane. Recording pipettes were pulled from filamented borosilicate glass pipettes (1.5 mm outer diameter, 1.10 mm inner diameter; Sutter Instruments) using a P-1000 Micropipette Puller (Sutter Instruments) and were filled with internal solution (in mM: 135 K-gluconate, 10 HEPES, 8 NaCl, 0.1 EGTA, 4 MgATP, 0.3 NaGTP, 10 Na-phosphocreatine; pH 7.2–7.3, 290–300 mmol/kg). For about half of our recordings, we prepared solutions to match the elevated osmolarity reported in zebra finch CSF (Bottjer, 2005). ACSF solutions were adjusted to 345–355 mmol/kg, and internal solutions to 335–345 mmol/kg. However, no noticeable differences in slice health or firing patterns were observed with this modification.

Electrodes had a resistance of 3–8 MΩ in the bath. Voltages were measured with a Multiclamp 700B amplifier (Molecular Devices Corporation) in current-clamp mode, low-pass filtered at 10 kHz, and digitized at 40 kHz with a Digidata 1440A. Pipette capacitance was neutralized, and 8–12 MΩ of series resistance was subtracted by bridge balance. Recorded voltage was corrected offline for measured liquid junction potential of 11.6 mV at 32 °C. Current injections and data collection were controlled by pClamp (version 10.4; Molecular Devices). Neurons were excluded from further analysis if the resting membrane potential was above –55 mV or if action potentials failed to cross –10 mV. Two hyperpolarizing current injections at different amplitudes (500–1000 ms) were used to monitor input and series resistance. Resting input resistance (*R*_*m*_), and capacitance (*C*_*m*_) were estimated from the responses to these step currents by fitting a sum of two exponential functions to the voltage decay (using Chebyshev polynomial regression). Recording sweeps were excluded if the input resistance, series resistance, or resting potential deviated by more than 20% from baseline.

Each neuron was stimulated with depolarizing step currents 2 s in duration at a range of amplitudes. Analysis of the responses followed the same procedures described previously (Chen and Meliza, 2018). Action potentials were detected as positive-going crossings of a threshold 35% of the difference between the peak of the first spike and its takeoff point. Only spikes that peaked at least 10 mV above the detection threshold and that were less than 8 ms in width (at half-height) were included. This procedure excluded the broad voltage oscillations observed in some neurons during prolonged depolarization. Cells were classified as narrow-spiking if the spike width was <0.8 ms or if maximum steady state spike rate was greater than 30 Hz, and were excluded from analysis.

The response duration in each sweep was measured as the time elapsed between the first and the last spikes of the pulse. In responses with only one spike, the duration (∆*t*) was defined as the width of the spike plus the duration of the afterhyperpolarization. The emphasis of this study was on phasic firing caused by a low-threshold potassium current, which produces outward rectification at around –60 mV (Chen and Meliza, 2018). We therefore excluded sweeps in which depolarization block occurred, as evidenced by a steady-state potential greater than –45 mV. However, sweeps with high steady-state potentials were not excluded from 5 neurons that had unusually high rheobases and had to be depolarized above –45 mV to spike at all. Cells were classified as phasic if they did not fire during the last 500 ms of the step current in response to any magnitude current; otherwise, they were classified as tonic. Note that in contrast to our previous study (Chen and Meliza, 2018), we no longer distinguish an “intermediate-spiking” class, because the immmunofluorescence data suggest that Kv1.1, which is likely to be the underlying low-threshold potassium channel that drives phasic firing, is distributed continuously across the population.

### Immunofluorescence

Animals were administered a lethal intramuscular injection of Euthasol and perfused transcardially with a 10 U/mL solution of sodium heparin in PBS (in mM: 10 Na2HPO4, 154 NaCl, pH 7.4) followed by 4% formaldehyde (in PBS). Brains were immediately removed from the skull, postfixed overnight in 4% formaldehyde at 4 °C, cryoprotected in 30% sucrose (in 100 mM Na2HPO4, pH 7.4), blocked saggitally into hemispheres, embedded in OCT, and stored at –80 °C. Sections were cut from one hemisphere at 40 µm on a cryostat, rinsed four times in PBS, and then incubated in citrate buffer (0.01 M, pH 8.5) at 80 °C for 25 minutes. The sections were cooled to room temperature, washed twice in PBS, permeabilized in PBS-T (0.1% Tween in PBS), and blocked in 5% goat serum and 2% glycine in PBS-T. The tissue was stained for Kv1.1 using a monoclonal mouse antibody (1:500 in blocking solution; UC Davis/NIH Neuromab clone K20/78; RRID:AB_10673165) for 60–72 h at 4 °C. Sections were washed 4 times for 15 min in PBS-T and then incubated with a fluorescent secondary antibody (1:2000 in PBS-T; goat anti-mouse IgG1 conjugated to Alexa Fluor 488; Invitrogen; RRID:AB_2534069). The tissue was washed twice with PBS, counterstained for Nissl (Neurotrace 640/660, 1:1000 in PBS; ThermoFisher, catalog N21483; RRID:AB_2572212), washed four times with PBS, and then mounted and coverslipped in Prolong Gold with DAPI (ThermoFisher, catalog P36934; RRID:SCR_015961).

Stained tissue was imaged with a 40X objective (water immersion, NA 1.2) on a Zeiss LSM 800 confocal microscope in stacks with 0.44 µm between optical sections. Higher-power images for illustration were obtained with a 63X objective (water immersion, NA 1.2). At least three stacks were imaged in each section in semi-standardized locations that spanned the dorsoventral and caudorostral axes. Images were collected from at least two sections in each animal, preferably one each from the medial and lateral subdivisions of CM. Left and right hemispheres were balanced across groups. To quantify expression, we counted the number of Kv1.1-positive neurons in each stack manually in Imaris (version 9.2.1) after using the automatic background subtraction, with the default smoothing radius of 39.9 µm. The total number of neurons in each stack was counted from the Nissl channel (in a few cases, we used DAPI when the Neurotrace staining was poor) using the automatic spot detection function in Imaris, followed by manual curation. Samples were processed and analyzed in three batches of 8–11 animals each, and each batch contained samples from every condition. Identical laser power and gain settings were used for the Kv1.1 channel in all the samples from each batch, but the Nissl and DAPI settings were adjusted freely to optimize signal quality. Experimenters were blind to animal identity and rearing condition throughout staining, image collection, and analysis, with the exception of one brain (CR, P19) that was repeated using the other hemisphere because none of the sections in the first batch contained CM.

The immunogen for the anti-Kv1.1 antibody was a synthetic peptide from the (intracellular) C-terminus of rat Kv1.1. The corresponding sequence in the zebra finch ortholog (KCNA1) is 74% identical, and a BLAST query against the zebra finch genome yielded no other hits. We validated the antibody by comparing staining patterns to *in situ* images from the ZEBrA gene expression brain atlas (ZEBrA Database, 2019). As in the atlas, cellular staining was much lower in L2a compared to the mesopallium and was absent in the habenula. In contrast, the two rabbit polyclonal antibodies we tried either failed to stain neurons in KCNA1-positive regions or stained cell bodies in KCNA1-negative regions. Each staining batch included control sections that were incubated in blocking solution without primary antibody. Staining from the secondary antibody in these sections was present in the neuropil but essentially nonexistent in cell bodies.

### Statistical analysis

Statistical inference was performed using generalized linear mixed-effects models (GLMMs) with bird and family as random effects to account for repeated measures. Age group and rearing condition were included as fixed effects. Contrasts were planned with P18–P20 and FR as baselines.

For average response duration, individual sweeps were treated as the unit of observation. The error model was Gaussian, and neuron was added as a random effect. The variance of this effect was allowed to differ by age and condition (i.e., we estimated the between-neuron variance separately for each age/condition, which is consistent with the obvious heteroskedasticity of the data). The rationale for not pooling all the duration measurements within neurons is that individual responses tended to be either sustained or transient, and the duration did not depend much on the stimulus amplitude. GLMMs provide partial pooling (Moen et al., 2016), which made the estimates less sensitive to outlier trials in which spontaneous activity caused an isolated spike to occur near the end of the current step. The credible intervals in this sweep-level model were in better agreement with the raw data.

The proportion of phasic neurons was estimated using a more stringent approach in which neurons were only classified as phasic if they did not fire any action potentials during the last 500 ms of the current step. In this model, individual neurons were the unit of analysis and the dependent variable had a binomial error model.

The proportion of Kv1.1-positive neurons was also estimated with a binomial GLMM, with image stacks treated as the unit of observation. Sex, rearing condition, and mediolateral position were included as fixed effects. Section, bird, family, and processing batch were included as random effects.

Because not every animal could be sexed and because dividing up the groups by sex reduced the effective sample size, the effect of sex on electrophysiological properties was analyzed separately in a model that only included CR animals. The electrophysiology rescue experiments were also analyzed in a separate model that only included animals between P30–P40.

Parameter estimates and credible intervals were obtained using MCMCglmm (version 3.5.2; Hadfield, 2010), which draws random samples from the joint posterior distribution of the model using Markov Chain Monte Carlo sampling. Prior distributions were weakly informative: for fixed effects, normal with variance of 1000 and for variances, an inverse Wishart with variance 1 and degree of belief 0.002. The number of samples and thinning were chosen to achieve adequate mixing (Gelman-Rubin diagnostic ≤ 1.01) and an effective sample size of at least 1000 for every parameter of interest. Statistical tests were performed by determining the proportion of samples that exceeded the value for the null hypothesis; for example, the average response duration for P18–P20 FR birds was greater than the value for P65–P70 FR in 960/1000 samples, leading to a p value of 0.04.

## Acknowledgments

We thank M. Bjoring for assistance analyzing acoustic recordings, F. Ahsan and K. Hess for help with analyzing immunofluorescence images, and G. Edgerton for feedback on the manuscript

## Author contributions

Conceptualization, ANC and CDM; Formal analysis, CDM; Funding acquisition, CDM; Investigation, ANC and CDM; Supervision, CDM; Visualization, ANC and CDM; Writing, ANC and CDM.

## Supplementary Figures

**Fig. S1.**
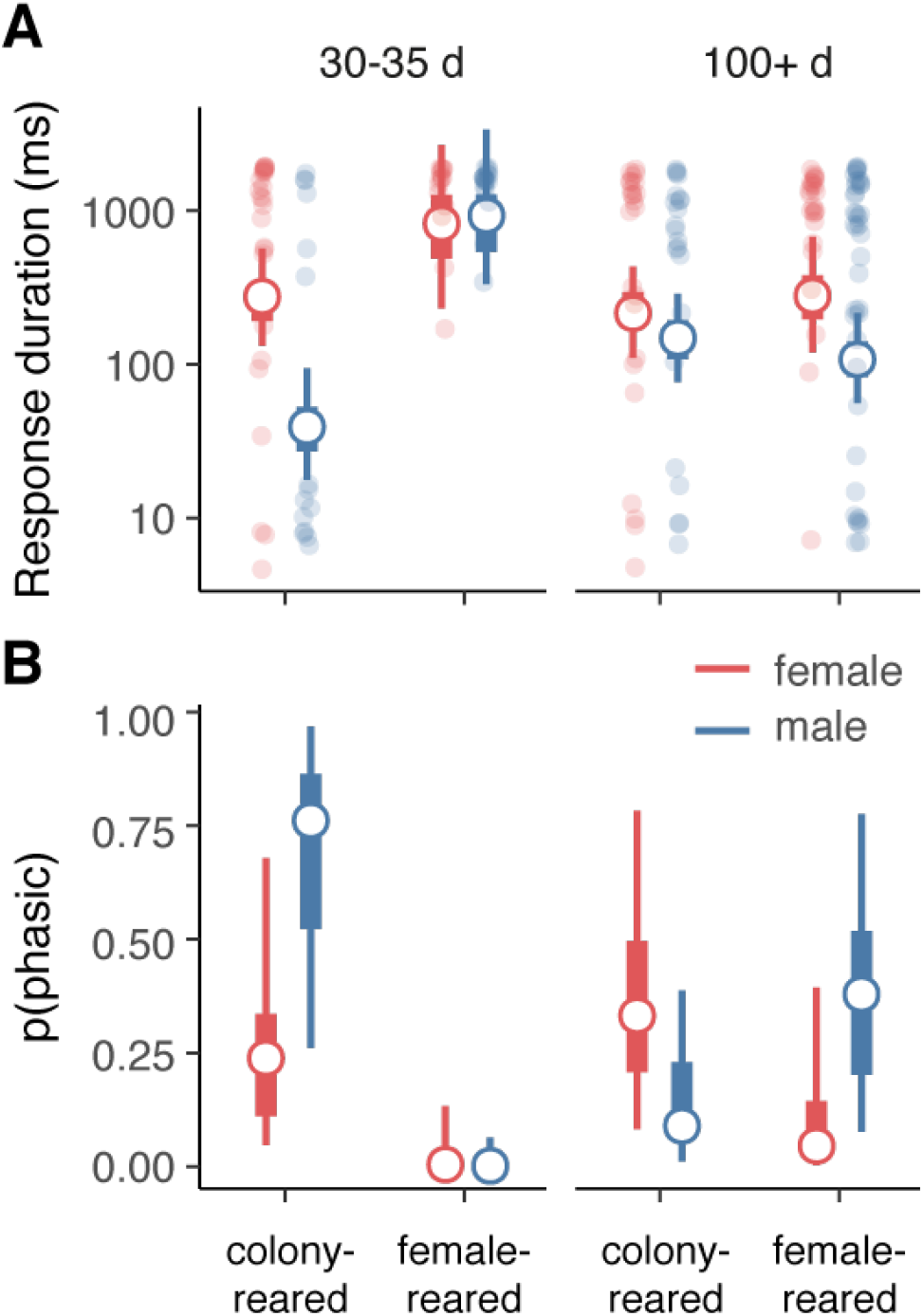
Effect of sex and rearing condition on phasic excitability during song memorization (P30– P35) and in adults (P100+). (**A**) In colony-reared juveniles, responses were shorter (more phasic) on average in males compared to females (GLMM: p = 0.004). In adults, there were no difference between colony-reared males and females (p = 0.51), but there was a trend for responses to be more phasic in female-reared males (p = 0.096). This could reflect the effects of self-generated song in the isolate males. The adult female-reared females were isolated from their male siblings at P40 so that they were not exposed to song of any kind. (**B**) A similar pattern is seen in the proportion of neurons that are phasic, but none of the differences are statistically significant using this more stringent binary criterion. As in main text figures, thick and thin whiskers indicate 50% and 90% credible intervals, respectively.

**Fig. S2.**
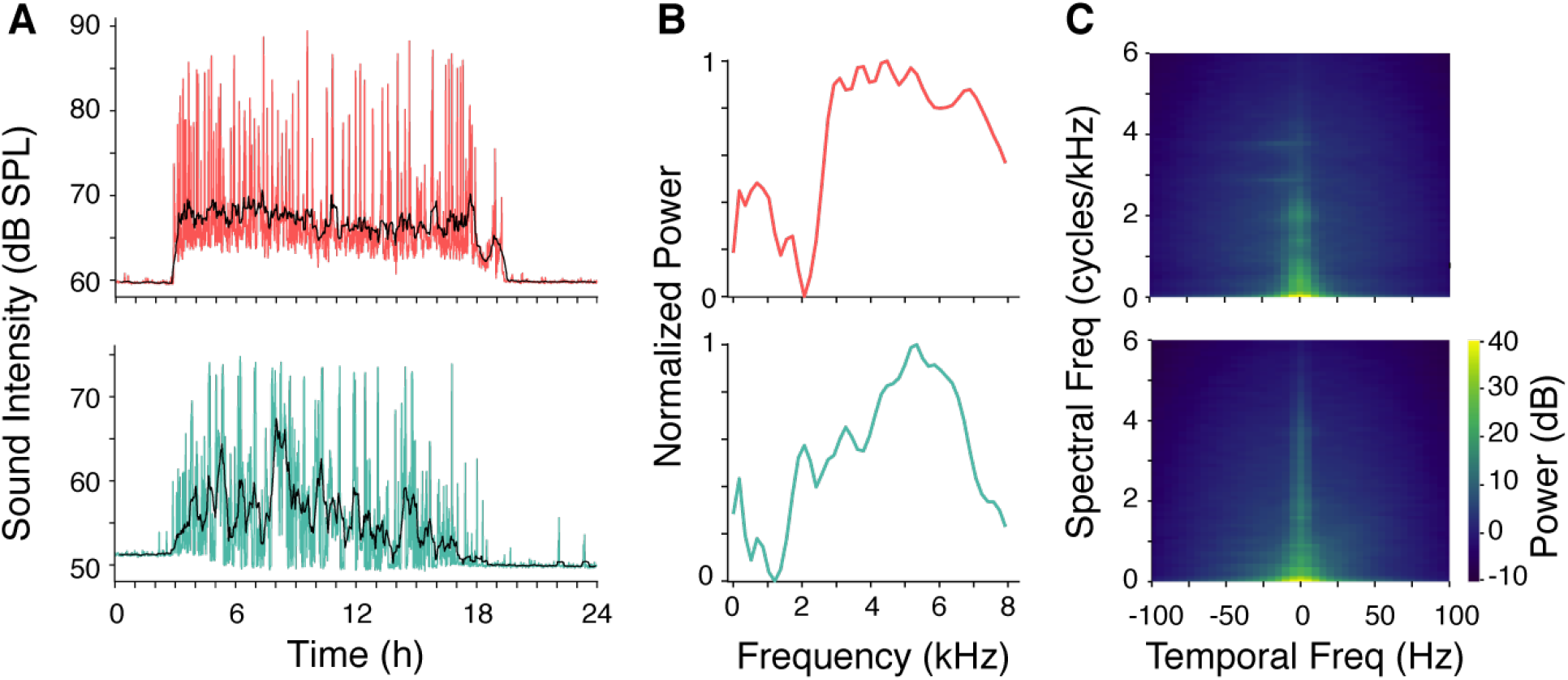
Example statistics of acoustic background obtained from nest box recordings of visual isolate (top row) and female-reared (bottom row) chicks. A small hole was drilled in the lid of the nest box, a piece of foam rubber padding with a cavity for a microphone was glued above the hole, and an omni-directional lavalier microphone (Shure SM93) was placed in the cavity. The signal was amplified and digitized at 44.1 kHz using a Focusrite Scarlett 2i2 and recorded directly to disk using custom software (jrecord version 2.1.4; https://github.com/melizalab/jill). Prior to installing the lid on the sound box, a recording was made of a 94 dB SPL calibration signal emitted by an R8090 (Reed Instruments) sound level calibrator placed immediately above the hole on the nest side of the lid. The gain on the preamp was kept constant throughout the recording. Recordings were obtained over several days while chicks were approximately a week old. (A) RMS amplitude of recording by time of day for a single exemplar day. Amplitude was quantified in non-overlapping intervals of 1 min (colored traces). Black traces show a moving average. (B) Average power spectra of recordings sampled during periods of relatively intense vocalization, normalized by maximum power. The dominance of power at high frequencies likely reflects begging calls (Elie and Theunissen, 2016). (C) Normalized modulation spectra (i.e., 2D FFTs of the log-scaled spectrogram) during the same intervals. In contrast to white noise, the noise experienced by chicks in both conditions is limited to relatively low-frequency spectral (0–3 cycles/kHz) and temporal (0–25 Hz) modulations. The modulation spectra are broadly similar between conditions, although there appears to be more power at higher spectral and spectrotemporal modulations in the colony.

